# Integrated analysis of single-cell transcriptome of liver cancer and cirrhosis reveals cell lineage similarity and prognostic-associated subpopulations

**DOI:** 10.1101/2022.11.03.515124

**Authors:** Mengsha Tong, Shijie Luo, Lin Gu, Zheyang Zhang, Chenyu Liang, Jingyi Tian, Huaqiang Huang, Yuxiang Lin, Jialiang Huang

## Abstract

**Background & Aims:** Liver cancer is one of the most leading causes of cancer deaths. Cirrhosis is an important risk factor for liver cancer, which is the result of over-fibrosis caused by diffuse and long-term liver damage. Despite extensive research, a systematic study for characterizing similarity between liver cancer and cirrhosis at single cell resolution is still lacking.

**Methods:** We established a data analysis framework to elucidate cell lineage similarity between liver cancer and cirrhosis to discover prognostic-associated subpopulations. We integrated single-cell transcriptome data from liver samples at normal, cirrhotic and cancer conditions, which totally contained 78,000 cells. Gene regulation analysis, cellular interactions and trajectory analysis were performed to characterize cirrhosis-like cell subpopulations. Bulk transcriptomes were used to discover prognostic-associated subpopulations.

**Results:** By aligning cellular subpopulations across different samples, we found remarkable similarity between *KNG1*^+^ hepatocytes in cirrhosis and *PGA5*^+^ hepatocytes in HCC. Furthermore, gene regulation analysis and cellular interactions implicated E2F1, FOXA2, EGF, CDH and ANGPTL signaling in maintaining cirrhosis-like ecosystem. Strikingly, subpopulations with higher expression of cirrhosis-like signatures were associated with patients’ worse survival.

**Conclusions:** We revealed a previously unexplored cirrhosis-like ecosystem of liver cancer, which could provide novel biomarkers for therapeutic interventions in HCC. Core analysis modules in this study were integrated into a user-friendly toolkit, SIM^scRNA^(https://github.com/xmuhuanglab/SIM-scRNA), which could facilitate the exploration of similarity and heterogeneity between precancerous diseases and solid tumors.

## 1. Introduction

Liver cancer is one of the most leading causes of cancer deaths globally(1). Clinical histopathological subtypes include hepatocellular carcinoma (HCC) and intrahepatic cholangiocarcinoma (iCCA)(2, 3). Liver diseases could be induced by dietary or viral factors, leading to fatty liver, cirrhosis, and eventually cancinomas(1). In previous studies, approximately 90% of patients with HCC had a history of cirrhosis(4). About 50% of patients with original sclerosing cholangitis developed bile duct carcinoma within 24 months(5). Lots of clinical reports revealed that cirrhosis is one of the most important risk factors for HCC and iCCA(6–13)(**Table 1**). Cirrhosis is the terminal stage of liver fibrosis, in the process of wound healing and scar tissue. It could affect hepatocytes, endothelial cells, hepatic stellate cells (HSCs) and a small number of monocytes. The main clinical manifestations of cirrhosis are impaired hepatocyte function, increased intrahepatic resistance and the development of liver cancer(14, 15).

**Table 1.**
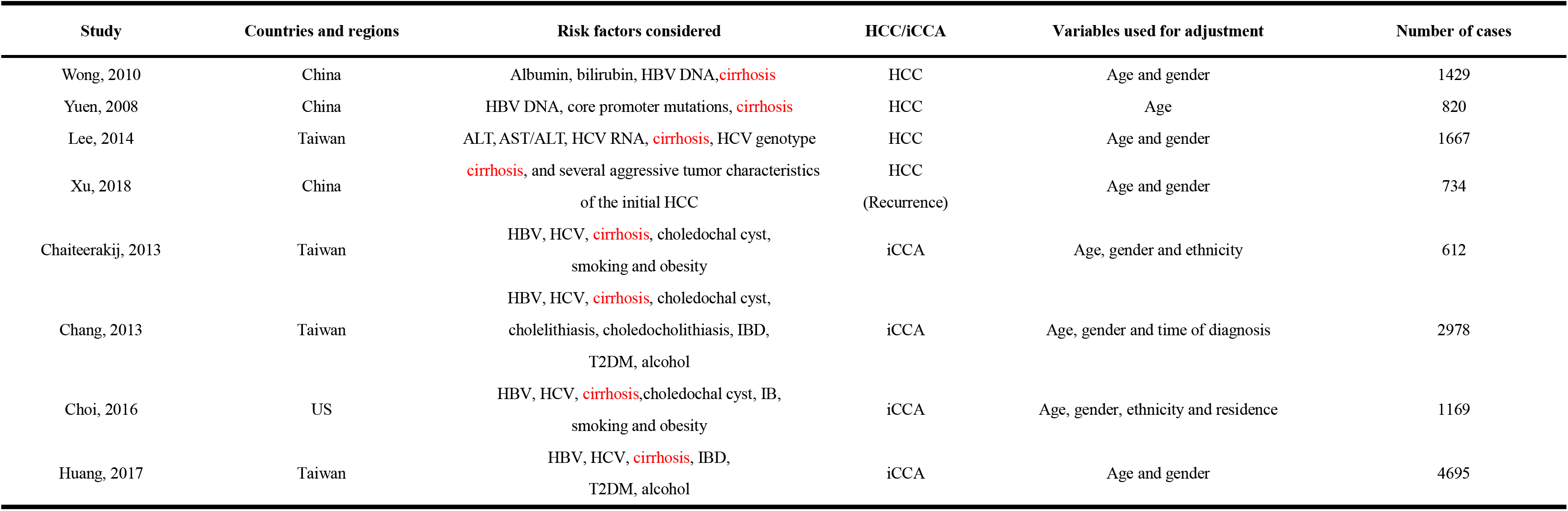
Summary of risk factors in liver cancer.

| Study | Countries and regions | Risk factors considered | HCC/iCCA | Variables used for adjustment | Number of cases |
| --- | --- | --- | --- | --- | --- |
| Wong, 2010 | China | Albumin, bilirubin, HBV DNA, **cirrhosis** | HCC | Age and gender | 1429 |
| Yuen, 2008 | China | HBV DNA, core promoter mutations, **cirrhosis** | HCC | Age | 820 |
| Lee, 2014 | Taiwan | ALT, AST/ALT, HCV RNA, **cirrhosis**, HCV genotype | HCC | Age and gender | 1667 |
| Xu, 2018 | China | **cirrhosis**, and several aggressive tumor characteristics of the initial HCC | HCC<br>(Recurrence) | Age and gender | 734 |
| Chaiteerakij, 2013 | Taiwan | HBV, HCV, **cirrhosis**, choledochal cyst, smoking and obesity | iCCA | Age, gender and ethnicity | 612 |
| Chang, 2013 | Taiwan | HBV, HCV, **cirrhosis**, choledochal cyst, cholelithiasis, choledocholithiasis, IBD, T2DM, alcohol | iCCA | Age, gender and time of diagnosis | 2978 |
| Choi, 2016 | US | HBV, HCV, **cirrhosis**, choledochal cyst, IB, smoking and obesity | iCCA | Age, gender, ethnicity and residence | 1169 |
| Huang, 2017 | Taiwan | HBV, HCV, **cirrhosis**, IBD, T2DM, alcohol | iCCA | Age and gender | 4695 |

Cirrhosis is commonly referred as “field cancerization” or “field effect” for it serves as the microenvironment that fosters initiation and promotion of neoplastic clones by facilitating genetic aberrations and cellular transformation. The carcinogenic ‘‘field effect’’ in the cirrhotic liver enables the exploration of liver cancer diagnostics and therapeutics(6). Maillyet al. revealed that fibrosis occurred during the transition from cirrhosis to HCC in a mouse induction model(6, 16, 17). However, the cellular and molecular connections and pathological mechanisms between liver cancer and cirrhosis remain unclear. Most of studies relied on bulk transcriptomes or statistical studies of clinical information (**Table 2**), which might partially obscure the characteristics of different cellular subpopulations and failed to quantify the similarities between liver cancer and cirrhosis at the cellular level. Single-cell sequencing technology could assist us in finding the molecular characteristics of microenvironment within cirrhotic and liver cancer samples. Another important issue should be noticed is that how to characterize similarity by integrating single-cell transcriptomic data of cancer samples from different sources or periods. Liu et al. performed alignment analysis of scRNA-seq data from colorectal cancer and the autologous liver metastasis to reveal cellular cross-talk signaling within the tumor microenvironment (18). Dong et al. compared single cell transcriptomes of human neuroblastoma (NB) and fetal adrenal glands to showed that malignant states could recapitulate the proliferation status of chromaffin cells in normal samples (19). Ankur Sharma et al. revealed that a novel onco-fetal ecosystem drived immunosuppression state in human fetal liver and HCC(20). These studies provide inspiration and technical supports to explore the subpopulations of cirrhosis-like cells that occur in liver cancer.

**Table 2.** Similar signaling pathways between cirrhosis and liver cancer.

| Study | Disease type | Biological process | Organism | Experiment |
| --- | --- | --- | --- | --- |
| Song S, 2022 | HCC | EGFR/MET stabilizes suspended HCC cells and avoids the killing of leukocytes by regulating the Ras/MAPK pathway | Human | Knockdown of EGFR/MET, inhibitor of EGFR/MET |
| Kim E, 2019 | HCC | β-catenin in the AJ complex supports HCC cell survival by promoting EGFR signaling during the earlier stages of HCC, and that its Wnt signaling associated role is restricted to late-stage HCC | Human, Mouse | Triple knockout hepatocellular carcinoma (TKO HCC) cells(control/shRNA) |
| Yu Y, 2020 | HCC | E2F1/DDX11 axis promote HCC cell proliferation and invasion through activating PI3K/AKT/mTOR pathway | Human | DDX11 Knockdown/Overexpression |
| Su Q, 2019 | HCC | HIF-1α/TGF-β feed-forward signaling; Hypoxia and TGF-β induced Smad and PI3K-AKT pathways and EMT in HCC cells | Human | TGF-β, CoCl<sub>2</sub> or 1% O<sub>2</sub> treatment |
| Lu S, 2022 | iCCA | NNMT promotes the proliferation and metastasis of iCCA in vitro and in vivo; NNMT promotes the Warburg effect by activating the EGFR-STAT3 pathway | Human, Mouse | NNMT Knockdown/Overexpression |
| Li F, 2020 | iCCA | MNX1-AS1 enhances ICC cell proliferation, migration, invasion, and angiogenic ability; MNX1-AS1 can facilitate ICC progression via MNX1-AS1/c-Myc and MAZ/MNX1/Ajuba/Hippo pathway | Human | MNX1-AS1 knockdown |
| Yamanaka T, 2020 | iCCA | Tumour fibrosis is caused by cancer-associated fibroblasts (CAFs), which are reported to play a key role in the ICC microenvironment, and promote malignant behaviour such as proliferation, invasion and metastasis; CAF-secreted cytokines, such as IL-6, IL-8, CCL2 and CXCL12, can increase the aggressiveness of co-cultured cancer cells | Human, Mouse | Co-implanted with CAFs;Antifibrotic drug nintedanib treatment |
| Moreno T, 2022 | Cirrhosis | Loss of FOCAD disrupts hepatocyte homeostasis and leads to cirrhosis; The increased secretion of CCL2, is expected to worsen the fibrogenic and inflammatory processes at play in FOCAD-mutant patients | Human, Zebrafish | CRISPR-mediated FOCAD-knockout |
| Yang YM, 2019 | Cirrhosis | HAS2(up-regulated by transforming growth factor-β) actively synthesizes HA in HSCs and that it promotes HSC activation and liver fibrosis through Notch1 | Human, Mouse | BDL- or CCl<sub>4</sub>-induced(fibrosis); HAS2 Knockdown/Overexpression |
| Sun J, 2022 | Cirrhosis | lncRNA SNHG1 silencing was observed to downregulate SMURF1 by upregulating miR-15a, diminishing ubiquitination of UVRAG to activate ATG5/Wnt5a axis, which accelerated HLC differentiation from BMSCs and repressed liver fibrosis to inhibit cirrhosis | Mouse | BMSCs derived from lncRNA SNHG1-silenced mice |
| BOULTER L,2012 | Cirrhosis | In human diseased livers and in mouse models of cholangitis, MFB initiates Notch signaling within HPCs through the expression of the Notch ligand Jagged1, promoting the differentiation of HPCs into cholangiocytes | Human | Drug injection induces abnormality in mice |
| Takuma T, 2018 | Fibrosis/Cimrhosis/HCC | WD/CC14 treatment induced ductular reaction and hepatocyte proliferation; CC14 treatment accelerates WD-induced steatohepatitis and fibrosis | Mouse | WD/CC14 treatment |

In this study, we comprehensively collected 78,000 single cells across human healthy, cirrhosis and liver cancer samples. Importantly, based on comparisons of transcriptional programs, gene regulatory networks and cellular interactions, we identified similar subpopulations between liver cancer and cirrhosis.

## 2. Methods

### 2.1. Datasets collection and preprocessing

In our study, we downloaded single-cell transcriptome data from Gene Expression Omnibus (GEO) for patients with HCC (n=22, GSE151530)(21), iCCA (n=12, GSE151530), cirrhosis (n=4, GSE136103) and healthy samples (n=5, GSE136103) (22). In addition, the bulk transcriptomes of HCC were downloaded from TCGA (n = 371), GSE14520 (n = 219), GSE20140 (n = 80), GSE76427 (n = 115) and LIRI (n = 228). The bulk transcriptome data for iCCA were downloaded from TCGA (n=40), EMBL (n=73), GSE107943 (n=30), GSE119336 (n=15), and GSE32879 (n=16), respectively. We normalized the matrix of TCGA data by *vst* method from DEseq2 package(23). Datasets detected by Affymetrix microarray were normalized by Robust Multi-array Average algorithm(24). The microarray datasets from EMBL source were converted into gene names by *getGEO* (https://bioconductor.org/packages/release/bioc/html/GEOquery.html). Clinical information including gender, age, disease stages, history of cirrhosis and survival time were summarized in **Supplementary Data 1**.

### 2.2. Cell clustering and annotation

Normalization of data, selection of highly-variable genes, dimension reduction and clustering of single-cell transcriptome data was performed using Scanpy, a scalable Python-based package(25). The main functions used in this study included *sc.tl.pca*, *sc.tl.umap* and *scanpy.tl.louvain* function. *FindIntegrationAnchors* function in Seurat was used to correct the batch effect among samples from different resources. Then, major immune cell types were annotated according to these markers: Hepatocyte (ALB/TF/TTR), Cholangiocyte (EPCAM/KRT19/CD24), T cell (CD3D/CD3E/CD3G), Mesenchyme (PDGFRB/ACTA2/COLIA1), Endothelial (PECAM1/CDH5/ICAM2), B cell (CD79A/CD79B/CD19), MPs (CD68/CD14/ITGAX), etc. Minor cell subpopulations were named according to the marker genes found by re-clustering (**Supplementary Data 2**). Finally, the accuracy of the annotated results was checked by using *sc.pl.correlation_matrix function*.

### 2.3. Sample preference of each subpopulation

To quantify the preference of each subpopulation, we calculated the observed and expected cell numbers of subpopulations in each sample(i) and apply chi-squared(ii) to test the distribution of subpopulations across samples deviates from random expectations. The subpopulations preference in a specific sample was defined according to Ro/e(26), and filtered out subpopulations with fewer than 40 cells that appeared simultaneously in healthy or cirrhotic samples.

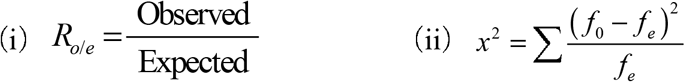

### 2.4. Identification of the Cirrhosis-like cell types

We applied several statistical methods to identify cell types in liver cancer samples capable of expressing features of cirrhosis. OriginAligner, Consensus non-negative matrix factorization (cNMF), PAGA connectivity and Jaccard similarity were used to calculate the similarity of two target subpopulations from cirrhotic and liver cancer samples. Gene regulatory network (GRN) correlation, and similar characteristics of cellular communication networks in different samples were also computed. Finally, the prognostic risk markers were screened from the similar features (**Figure 1a**). Core analysis modules were integrated into a user-friendly toolkit, SIM^scRNA^ (https://github.com/xmuhuanglab/SIM-scRNA)

**Figure 1.**
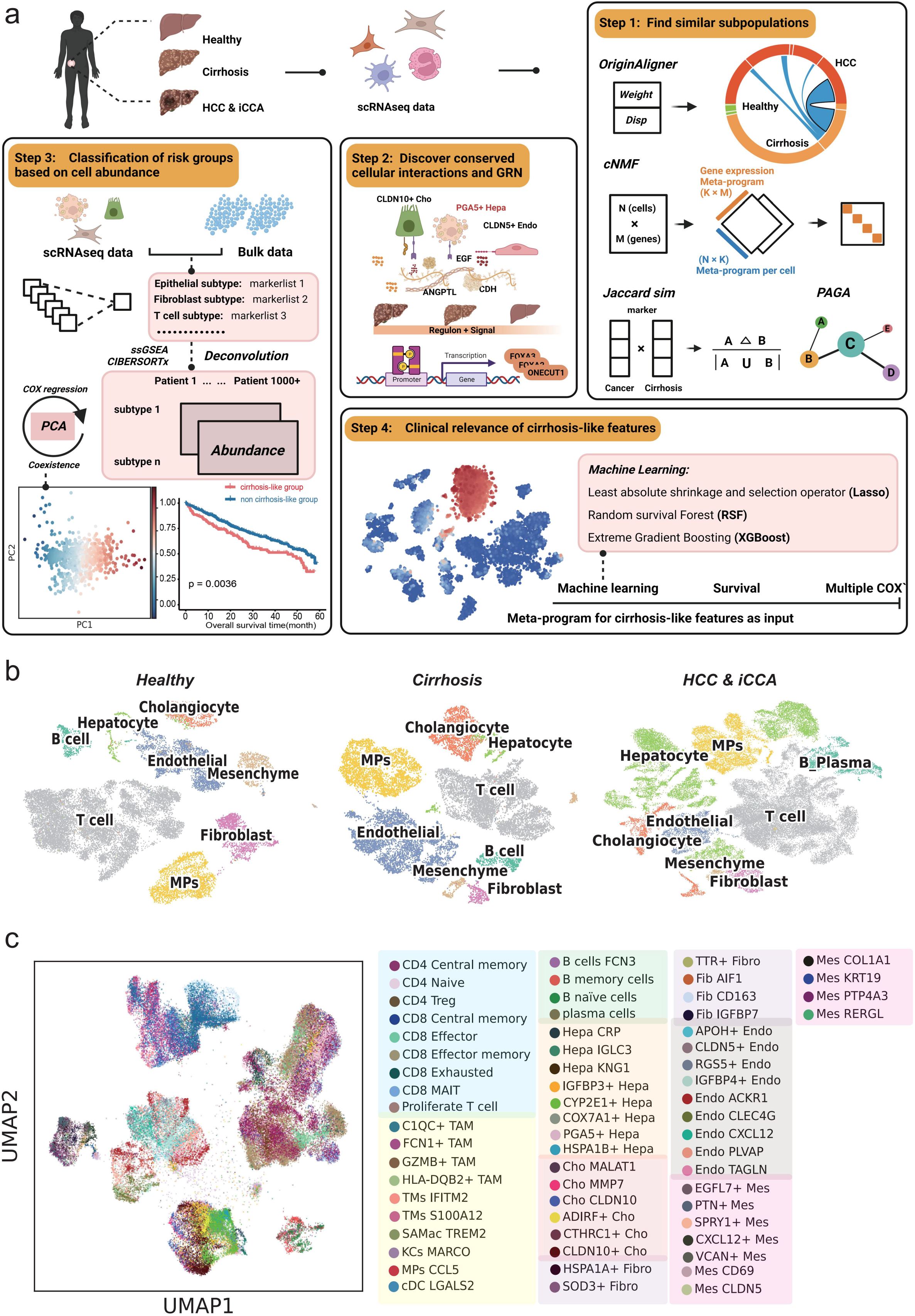
The framework and cell annotations in this study. **(a)** The workflow of our study: Single-cell and bulk transcriptomes from healthy, cirrhosis and liver cancer samples, were collected. OriginAligner, cNMF, PAGA connectivity and Jaccard similarity were used to identify similar subpopulations (Step1). This framework also used GRN and cellular interaction to characterize the impact of cirrhosis-like features on the tumor microenvironment (Step2). Deconvolution were used to construct the hierarchy of patients and demonstrate the prognostic value of cirrhosis-like signatures (Step3). Finally, machine learning was used to filter clinically relevant cirrhosis-like signatures (Step4). The workflow was drawn at https://biorender.com. **(b)** Uniform manifold approximation and projection (UMAP) plot showing three different sample sources of cell types. **(c)** UMAP plot showing the 63 subpopulations of all samples.

#### 2.4.1. OriginAligner

Here we designed a modified statistical method OriginAligner based on the hypothesis that cells resemble transcriptome characteristics in different samples might have similar microenvironments or niches(27). We firstly normalized the raw count expression matrix, then screened the genes with coefficient of variation ranked at top 500 by calculating sd/mean. The top 500 genes with high specificity for all subpopulations were further screened using Entropy Weighting method. After that, a query-reference framework based on k-nearest neighbors (kNN) was employed: tumor subpopulations were used as reference objects and healthy/cirrhosis subpopulation were used as query. A phenotypic marker index for each query type was defined by calculating the proportion of query cells labeled with the corresponding sample type (**Figure 2a**).

**Figure 2.**
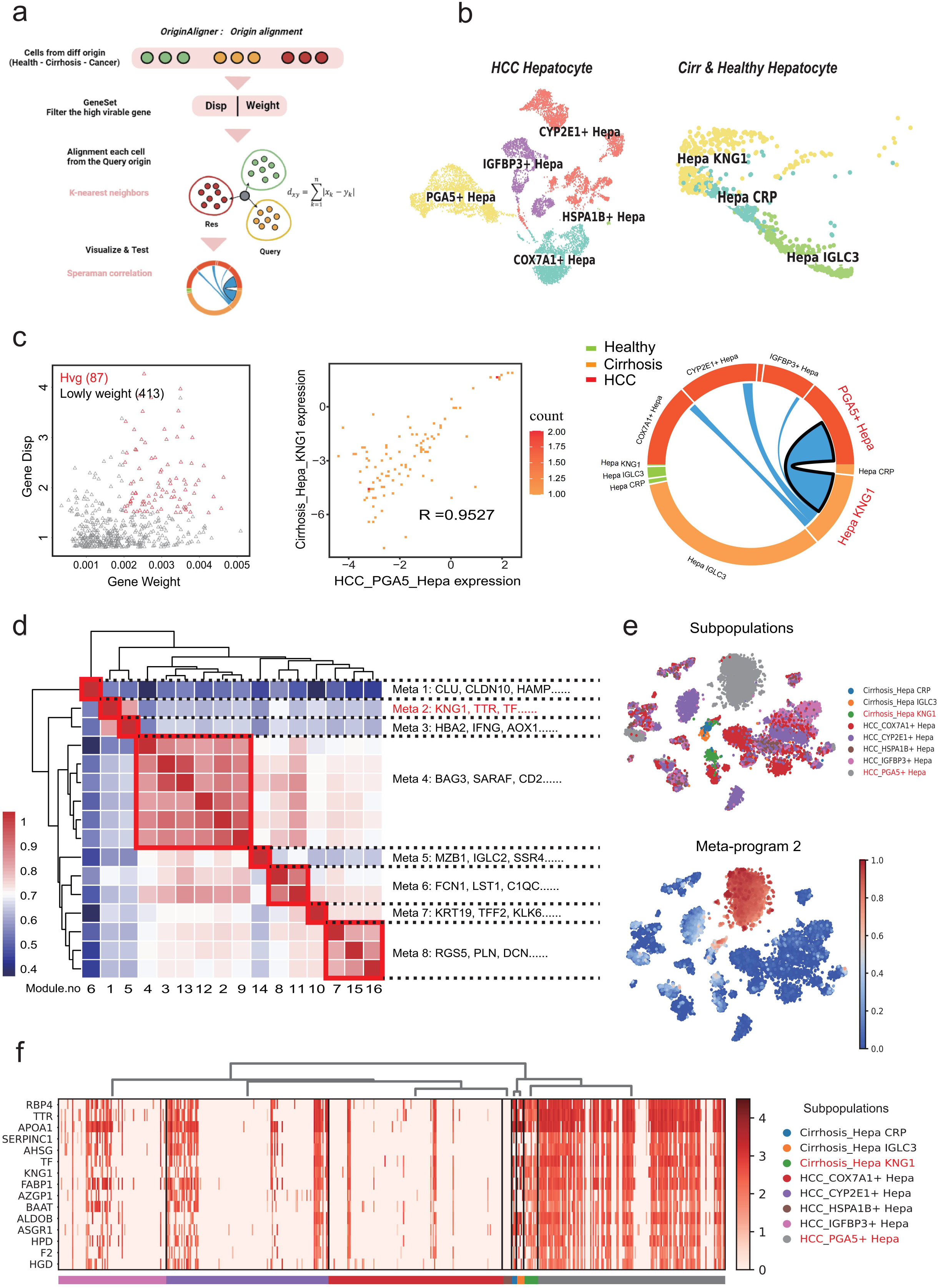
Identification of Cirrhosis-like cell subpopulations. **(a)** Schematic diagram of the OriginAligner method: Cells from cirrhosis and healthy sample were aligned to cells from liver cancer (HCC or iCCA) samples (reference cells) according to their molecular similarities based on KNN (measured by distance within the highly variable genes which filtered by dispersion and weight). **(b)** UMAP visualization of hepatocytes colored by subpopulations among different samples. **(c)** Genes screened by the dispersion of genes and the weighting of each hepatocyte cell subset (**left**). Correlation of highly variable genes expression in two similar subpopulations were obtained by screening, with each point representing a gene (**middle**). Circle plot showing the molecular similarities of various subpopulations of hepatocytes in three tissues (**right**). **(d)** Heatmap showing correlation of all programs derived from cNMF analysis of HCC and cirrhosis (k=16). Eight highly correlated meta-programs were highlighted. **(e)** t-SNE visualization of hepatocytes colored by subpopulations and meta-program 2. **(f)** Heatmap showing top 15 genes commonly expressed in meta-program 2.

#### 2.4.2. Connectivity score

We calculated connectivity scores using *scanpy.tl.paga* function to quantify the similarity of subpopulations between cirrhosis and liver cancer. PAGA connectivity(28) was the ratio of the number of inter-edges between the clusters to the number of expected inter-edges under the random assignment(i). The connectivity score indicates the confidence level that actual connectivity exists, which allows for discarding spurious and noise-related connectivity. *Cij* represented connectivity between cell subpopulation *i* and cell subpopulation *j*. *Eobs* represented the number of observed edges between cell subpopulations. The number of edges expected (*Eexp)* was the number of edges (*ei*) of the cell subpopulation *i* and all other cell subpopulations multiplied by the number of cells contained in cell subpopulation *j*. The corresponding *ej* and *ni* were added to devide by *n (total number of cells)-1* (ii), and the larger calculated value, the greater similarity between the two subpopulations.

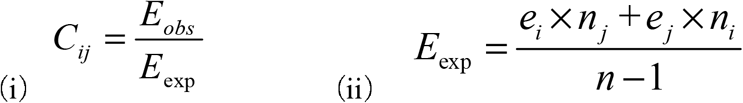

#### 2.4.3. Consensus non-negative matrix factorization

We dissected gene expression programs based on non-negative matrix factorization (NMF), which were repeated 100 times for each cell type and a set of consensus programs were computed by aggregating results from all 100 runs. The cNMF v1.4 python package was used to perform the analysis of consensus NMF (29). We determined the optimal number of programs (p) for each cell type by maximizing stability and minimizing error of the cNMF solution as well as ensuring the programs were biologically coherent.

#### 2.4.4. Jaccard similarity

Transcriptional similarity between two subpopulations were also evaluated by the Jaccard similarity coefficient. After filtering out subpopulations with similar transcriptome features in cirrhotic and liver cancer samples by OriginAligner and PAGA, we further selected marker genes using *scanpy.tl.rank_genes_groups* function. We selected top 60/100/200 marker genes according to the characteristics of subpopulations of different scales. A and B represented marker gene lists of subpopulations.

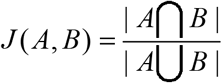

### 2.5. Regulon inference and signature enrichment analysis

#### 2.5.1. pySCENIC

Single-cell regulatory network inference and clustering(SCENIC)(30) from the python package ‘pySCENIC’ was used to explore similar characteristics at the gene network regulatory level. The potential transcription factors were found by GENIE3 using co-expression between genes. The high-frequency motifs appearing near the gene transcription initiation site were used. The motifs (NES>3) found in the database were compared to calculate the regulators enriched by the target gene. Finally, based on the expression value of the gene, the specific regulator activity specific to each cell was calculated with AUCell function. To obtain the correlation between each subpopulation, Spearman correlation was used.

#### 2.5.2. Palantir

Palantir(31) was used to model the pseudotime trajectory of hepatocytes between liver cancer and cirrhosis. The calculated pseudo-time assigned a relative distance to the initial cell for each cell, and the calculation process was as follows: The filtered gene expression matrix was normalized using *sc.pp.normalize_per_cell* function. Highly variable genes were selected for Principal component analysis (PCA). Clustering analysis was performed using *palantir.utils.run_diffusion_maps* function. Thus, we could set a random cells from healthy subpopulation as a starting point, then used the random cells corresponding subpopulations of cirrhosis and liver cancer to calculate the migration probability. For each sample, we evaluated the results from pySCENIC, as well as proven TFs that might promote liver cancer progression, to observe two branches of the cell lineage corresponding to cirrhosis-like cell subtypes.

### 2.6. Cell Cell Communications

NicheNet(32)(https://github.com/saeyslab/nichenetr) was used to investigate cell to cell interaction. This method uses public databases (ENCODE, KEGG, PhoshoSite) to map the targets for transcription factors and receptors for the provided dataset. In this study, receiver cell type was specified as hepatocytes and sender cell types were T cell and Endothelial populations. The condition of interest were ‘HCC’ and ‘Cirrhosis’ as compared to ‘Healthy’ region. CellChat(33) was used for analyzing signaling pathways based on cell-to-cell interactions. The ligand and receptor information was from *CellChatDB.human*. Normalized expression profiles was required as input for CellChat analysis, and objects were created via *createCellChat*. In our analysis process, we identified the specific ligand-rcceptor (L-R) pairs of the target cell subpopulations in the cancer sample by *identifyOverExpressedGenes* function, based on the ligands with high expression in the target cell subpopulations. Function *netAnalysis_signalingRole_heatmap* was used to calculate the degree of interaction with other subpopulations in healthy, cirrhosis and liver cancer, respectively. We also used *netVisual_Bubble* function to plot some of the interactions of interest (L-R pairs).

### 2.7. Revealing prognostic value

#### 2.7.1. Deconvolution of gene expression

Normalized gene expression matrix from 33,686 single cells, belonging to 21 HCC subpopulations were used for GSVA(34) function and CIBERSORTx(35). Deconvolution was performed on bulk transcriptome, including 1,013 patient samples from 5 cohorts (TCGA, GSE14520, GSE20140, GSE76427 and LIRI). The abundance of the 21 HCC subpopulations were normalized to a sum of 1. The score represents the estimated proportion for each cell type. The normalized HCC cell type compositions of these patients were used for the PCA analysis. Euclidean distance was used to detect neighbors. We project the query dataset onto the PCs of the reference dataset and assign cluster labels.

#### 2.7.2. Glmnet COX regression

To identify the prognosis-related groups, we performed Cox regression analysis on 5 cohorts using an L1(LASSO) penalty with partial likelihood deviance loss. The optimal lambda value was determined based on Leave-one-out cross-validation. Coefficients for each HCC subpopulation were used to calculate clinical feature importance. Pearson correlations were calculated between the absolute abundance of each subpopulation with each clinical feature.

#### 2.7.3. CNV Estimation

We used infercnvpy(36) to infer CNVs from each cell type. We randomly selected 100 cells from each cell type in healthy samples as reference cells. We used log2 (TPM+1) to transform the expression profiles, and we used a cutoff of 0.1 for the minimum average read counts per gene among reference cells. All genes were sorted by start position and chromosome number. The chromosomal expression patterns were estimated from the averages of 100 genes as the window size and adjusted as centered values across genes.

#### 2.7.4. Risk prognosis signature

The gene sets with high survival risk factors were inferred from 5 cohorts, and the risk coefficient(37) were calculated for all genes using *cox.zph*. Risk factors were considered if Hazard_Ratio > 1, while Hazard_Ratio < 1 was considered as protective factor. Prognostic risk enrichment score, which means the enrichment of these gene sets in various cell types within healthy, cirrhosis and liver cancer. Prognostic risk enrichment scores and CNV score were used to screen for subpopulations that were under mutational stress and associated with high risk of survival status. To identify shared cirrhosis-like signatures associated with survival, we selected top 100 shared gene from cNMF as input for predicting. We constructed 3 machine learning models, including extreme gradient boosting (XGBoost, https://xgboost.readthedocs.io/en/stable/index.html), Lasso regression (glmnet, https://cran.r-project.org/web/packages/glmnet/index.html), and randomforest (https://cran.r-project.org/web/packages/randomForest/index.html). The AUC value of the Receiver Operating Characteristic (ROC) curve (https://rdocumentation.org/packages/ROCR/versions/1.0-11) was calculated to evaluate the performance of the prognosis prediction models.

To further investigate the correlation between cirrhosis-like features and patient prognosis, we defined cirrhosis-like hub gene within the target subpopulations. GSVA was used to calculate the enrichment scores of these features in all patients. Survival analysis was done using R package Functions *survminer* and *survival. surv_cutpoint* was used to divide the patients into high and low groups based on enrichment scores. The survival curve was estimated using the Kaplan-Meier method, and the log rank sum test was used for comparing difference of survival time. The independence of prognostic variables was analyzed using a multivariate Cox regression model, which was presented in the forest plot.

### 3. Results

### 3.1. Cell composition in Healthy liver, Cirrhosis, HCC and iCCA

We collected single-cell transcriptome data from 5 healthy livers, 4 cirrhosis samples, 22 HCC samples, and 12 iCCA samples with 22,853, 16,522, 34,287 and 4,934 cells, respectively (**Figure S1a**). First, we employed lineage specific markers to annotate 8 major cell types for each sample source (**Figure 1b**, **Figure S1b-S1d, Supplementary Data 2**). Next, we subdivided major clusters into subpopulations by re-clustering of these cells. 31 subpopulations were identified in primary liver cancer and 37 subpopulations in healthy and cirrhotic samples, respectively (**Figure 1c, Supplementary Data 2**). For example, in the annotation of each subpopulation of HCC and iCCA, hepatocyte could be divided into five subpopulations: *CYP2E1*^+^ Hepatocyte, *IGFBP3*^+^ Hepatocyte, *PGA5*^+^ Hepatocyte, *HSPA1B*^+^ Hepatocyte and *COX7A1*^+^ Hepatocyte. And hepatocyte within cirrhosis could be identified three clusters: Hepa KNG1, Hepa IGLC3 and Hepa CRP (**Figure S1c-S1e**, See Method). In different samples, each cell type have different enrichment preferences, we observed that hepatocytes are significantly enriched in HCC and cholangiocytes are significantly enriched in iCCA (p < 0.001, Wilcoxon test) (**Figure S1d**, **Figure S1e**). In addition, there were no significant differences for other cell types. Overall, this integrated atlas of 78,596 cells from healthy, cirrhosis and tumor tissue of the human liver enables to compare molecular connections of liver cancer and cirrhosis.

### 3.2. Discovering cirrhosis-like cell-types in primary liver cancer

To quantitatively characterize cirrhosis-like cell types within liver cancer and their potential features, we analyzed integrated atlas described above with a modified OriginAligner method (see Methods). It applied the kNN-based algorithm to align the cellular phenotypes across samples from different tissues. We used single cells from primary liver cancer as a reference and aligned each cell from healthy and cirrhotic to identify cell subpopulations of their nearest sample (**Figure 2a**). The results showed that epithelial cell subpopulations in primary liver cancer and cirrhotic samples exhibit a higher similarity compared to healthy samples. Additionally, since the relatively independent presence of cholangiocytes and hepatocytes in the two primary liver cancer datasets, the epithelial subpopulations of these two cancer types needed to be distinguished and annotated separately (**Figure 2b**, **Figure S2a-S2b**). OriginAligner found similar characteristics of *KNG1*^+^ hepatocytes in cirrhosis and *PGA5*^+^ hepatocytes in HCC. We considered that high dispersion genes might be influenced by the number of cells in the subpopulation, so we additionally calculated the weight of each highly variable gene for the subpopulationand 87 genes were screened (See Method). *KNG1*^+^ Hepatocyte in Cirrhosis and *PGA5*^+^ Hepatocyte in HCC have similar characteristics (**Figure 2c**). PGA5 mainly involved in K[RAS signaling pathway, bile acid metabolism, mitotic G2 M phase, mTOR pathway and DNA repair(38). In contrast, *KNG1*^+^ hepatocytes in healthy samples were not significantly associated with these two subpopulations (**Figure 2c**). This finding was consistent with the notion presented by the low-dose diethylnitrosamine (DEN)-induced cirrhosis-driven HCC rat model, which elucidated that the pattern of gene module dysregulation was comparable(39). They found the cross-talks between cell types inducted by growth signaling and stress response/ECM were relatively stronger in hepatocytes(39).

Furthermore, we identified recurrent expression programs from hepatocyte within cirrhosis and HCC using cNMF(See Method). We determined the number of cNMF programs and focused on those shared between *PGA5*^+^ hepatocyte and *KNG1*^+^ hepatocyte(**Figure S3a**).We obtained 8 malignant cell programs and found that malignant meta-programs 2 had the most similar molecular characteristics between *KNG1*^+^ hepatocyte and *PGA5*^+^ hepatocyte. These results validated that tumor cells might be closer to *KNG1*^+^ hepatocyte or *PGA5*^+^ hepatocyte and approximately 15 genes were expressed only in these two subpopulations (**Figure 2d-2f, Figure S3b**). We selected the intersection of top 100 genes from these two subpopulations and each meta-program revealed that cirrhostic cells might be closer to *PGA5*^+^ hepatocyte (**Figure 3a**). Besides, the *KNG1*^+^ signature gene enriched in the HCC-specific gene set, likewise, the *PGA5*^+^ signature enriched in the cirrhosis-specific gene set (**Figure 3b**). In addition, we screened 54 genes from subpopulations of cholangiocyte as input to OriginAligner (See Method). We found strong transcriptomic similarity between *CLDN10*^+^ cholangiocyte in cirrhosis and *CLDN10*^+^ cholangiocyte in iCCA (**Figure S2c**, **Figure S2d**), which was further verified using the Connectivity score and Jaccard similarity.

**Figure 3.**
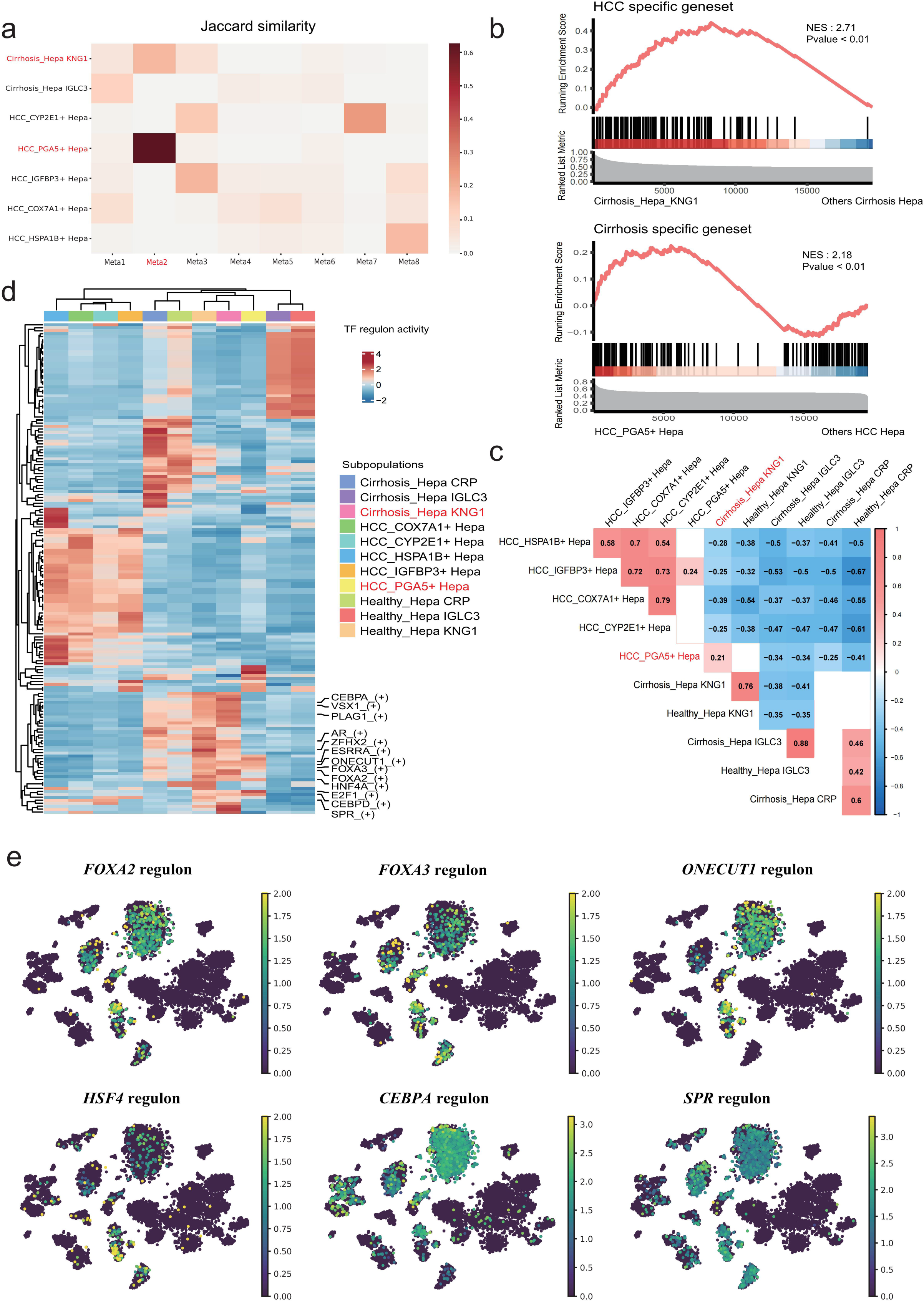
Gene regulatory network in Hepatocyte subpopulations. **(a)** Jaccard similarities of eight meta programs (x axis) with the signatures of seven subpopulations (y axis). **(b)** GSEA analysis for HCC and cirrhosis-specific gene sets. Genes were ranked by logarithmic fold change in the mean expression values of *KNG1*^+^ hepatocyte and *PGA5*^+^ hepatocyte, respectively. **(c)** Correlation of binary regulon activity percentage between each hepatocyte subpopulations. **(d)** Heatmap showing the activity of the top 15 SCENIC-inferred master regulons of seven subpopulations of hepatocyte. **(e)** t-SNE visualization of the SCENIC-regulon activity of six regulons (FOXA2, FOXA3, ONECUT1, HSF4, CEBPA, and SPR) in Hepatocyte subpopulations shown in **Figure 2e**.

In addition, during the annotation of cholangiocytes in iCCA, we found that SPP1 was specifically expressed in *CLDN10*^+^ cholangiocyte subtype. Current studies have shown that expression level of SPP1 in malignant glioma, liver cancer, bladder cancer and cervical cancer were closely associated with tumorigenesis, progression, and poor prognosis. In addition, the high expression status of this gene might serve as a new biomarker for early diagnosis or progression, invasion and metastasis(40) (**Figure S2d, Figure S2e**). Similarly, the results calculated from connectivity score found that high similarity between *TAGLN*^+^ endothelial, *TREM2*^+^ SAMac in cirrhosis, *CLDN5*^+^ endothelial, *FCN*^+^ TAM in HCC (**Figure S2f**). Scar-associated *TREM2^+^ CD9*^+^ subpopulations expanded in hepatic failure, differentiated from circulating monocytes, and were the major cell subpopulations that promoted liver fibrosis. Notably, *FCN*^+^ monocytes in colon cancer were highly similar to *CD14*^+^ monocyte in blood, possibly representing the monocyte populations would migrate into tumor with a tumor-specific transcriptional program. And *FCN1*^+^ monocytes could differentiate into *C1QC*^+^ TAM and *SPP1*^+^ TAM in different tumor microenvironments(18).

### 3.3. Conserved Gene Regulatory Network in Cirrhosis and HCC Hepatocytes

Our results from the OriginAligner, cNMF and PAGA analysis indicated a similar transcriptional program between cirrhosis and liver cancer. We hypothesized that the GRNs in these similar cell subpopulations could help to detect conserved transcription factors that might promote the transformation of cirrhosis to liver cancer. We assessed the correlation analysis among all identified regulators from hepatocyte in three samples and only *PGA5*^+^ hepatocytes in HCC and *KNG1*^+^ hepatocytes in cirrhosis showed a significant correlation (**Figure 3c**). Among the top15 regulatory factors selected in each hepatocyte subpopulation, *PGA5*^+^ hepatocytes in HCC and *KNG1*^+^ hepatocytes in cirrhosis had similar regulatory network, such as HSF4, FOXA2, ONECUT1, FOXA3, etc. (**Figure 3d, Figure 3e, Figure S4a**).

To further characterize whether these regulators can regulate both cirrhosis and liver cancer, or which potential regulators can determine the evolution of cirrhosis to liver cancer, we used Palantir to predict the transition from *KNG1*^+^ hepatocytes in cirrhosis to *PGA5*^+^ hepatocytes in HCC against the background of *KNG1*^+^ hepatocytes in healthy (**Figure S4b**, **see Method**). We found that HCC and cirrhosis shared already known liver cancer-specific regulators. According to previous reports, ONECUT1 and ONECUT2 had a superfluous role in the differentiation of hepatocytes or cholangiocytes. ONECUT1 stimulated Hnf1β expression to promote intrahepatic bile duct morphogenesis(41). Besides, previous studies had shown that FOXA2 inhibitd transcription of miR-103a-3p to induce upregulation of GREM2, thereby increasing LATS2 activity and phosphorylation of YAP (Hippo signaling pathway) to inhibit the migration and invasion of liver cancer cells(42), indicating that the regulatory role of FOXA2 might be significantly downregulated in HCC. TF MTDH mainly regulated the transition from hepatitis to HCC, BACH1 was responsible for promoting HCC growth, and HNF1A increases the tumor mutations burden. These regulators have conserved regulatory patterns in both subpopulations, suggesting that TFs that promote malignant proliferation might determining whether cirrhosis can transition to liver cancer. Meanwhile, we also identified some co-regulators in iCCA (**Figure S4c-S4f**), such as FOXA3, MEOX1 and NFE2L1. FOXA3 was highly expressed in intrahepatic cholangiocarcinoma tissues (61%)(43). The regulation of P311 expression by Meox1 contributed to promote the fibrosis process(44). In addition, ROS could cause immune escape of tumor cell through activation of NFE2L2-PDL1 pathway, and enhance tumor immune response through regulation of the NFE2L1-41BBL pathway(45).

Taken together, SCENIC analyses showed a similar transcriptional regulatory network associated with *KNG1*^+^ hepatocyte within cirrhosis.

### 3.4. PGA5^+^ Hepatocytes Exhibit Pro-inflammation and Pro- proliferation Interactions

Next, in order to characterize the similarities at cellular communication levels. We performed cell-cell interaction analysis using CellChat (**see Method**) to reveal ligand-receptor interactions in health, cirrhosis, and liver cancer. We compared the number of interactions and found that from health, cirrhosis to liver cancer, *PGA5^+^/KNG1*^+^ hepatocytes exhibited increasing interactions with immune cells in their own microenvironment (**Figure S5a**), especially the T cell subpopulations such as *CD4*^+^ Naïve, *CD4*^+^ Central memory, *CD4*^+^ Treg, proliferate T cell, etc. To demonstrate this finding, we calculated the correlation of the abundance of all subpopulations in five HCC bulk transcriptome data by using CIBERSORTx. *PGA5*^+^ hepatocytes were found to co-exist with *RGS5*^+^ endothelial, *CD4*^+^ naive T cell and *CD4*^+^ Treg in multiple data sets (**Figure S5b**). Next, we performed NicheNet (**see Method**) analysis to predict cellular interactions. Interestingly, NicheNet predicted that T cell-derived LTB could activate LTBR in HCC (**Figure S5c**), and LTB-LTBR was known to specifically expressed in liver cancer(46).

In addition, we selected the signals specifically expressed in the *PGA5*^+^ hepatocytes within HCC, such as ANGPTL, EGF, and CDH (**Figure 4a**). We found that *CD4*^+^ central memory, *CD4*^+^ Treg, and *CLDN5*^+^ endothelial and *PGA5*^+^ hepatocytes had closer contact (**Figure 4b**). ANGPTL was not only angiogenic factors, but also protein with multiple functions in carcinogenesis, cancer growth, proliferation, invasion, and metastasis(47). Notably, recent reports have demonstrated overexpression of EGFR resulted in strong expression of many oncogenes(48). CDH was secreted by epithelial cells and played a decisive role in the ferroptosis of tumor cells(49). In these three signaling contexts, *KNG1*^+^ (healthy/cirrhosis) / *PGA5*^+^ (HCC) hepatocyte played an increasingly important role in microenvironmental cellular communication (**Figure 4c**). Meanwhile, we found the same pattern in intrahepatic cholangiocarcinoma (**Figure S5d-S5e**). Taken together, these results further demonstrated the nexus between *PGA5*^+^ hepatocyte in HCC and *KNG1*^+^ hepatocyte in cirrhosis within the tumorigenesis ecosystem.

**Figure 4.**
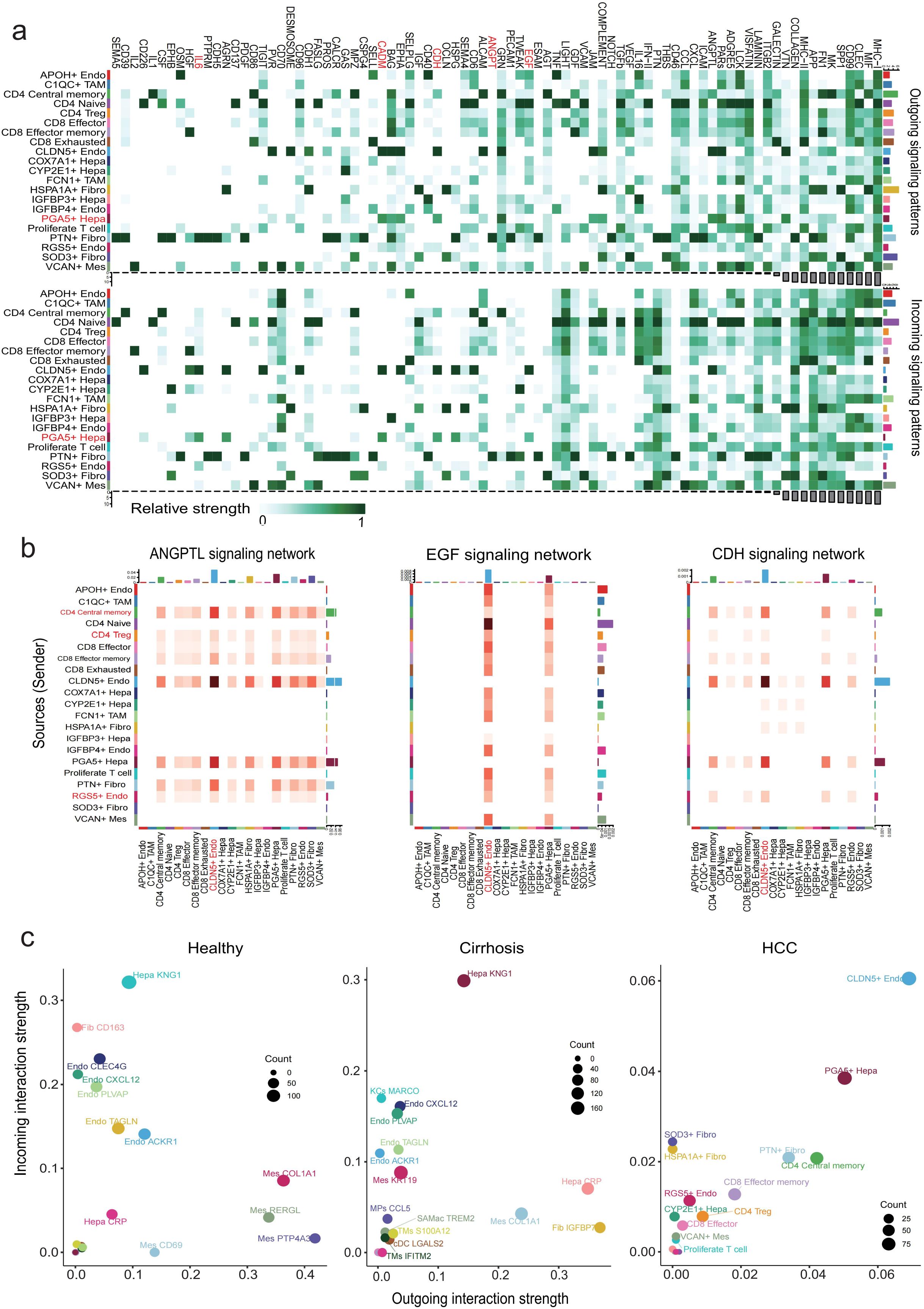
Characterization of cirrhosis-like subpopulations through cellular communication. **(a)** The heatmap shows the expression intensity of all signals of seed (**top**) and reverser (**bottom**) in the HCC subpopulations. **(b)** Heatmap of ANGPTL, EGF, CDH signaling pathway between sender cell types (y axis) and receiver cell types (x axis). Color reflects communication strength. **(c)** The outgoing interaction strength and incoming interaction strength of each cell type in different sample origins. Dot size is proportional to the number of inferred links (both outgoing and incoming) associated with each cell group.

### 3.5. Hierarchy composition of PGA5^+^ hepatocytes is associated with poor prognosis

We next sought to investigate the functional, biological and clinical properties of HCC cell subpopulations. Firstly, we performed gene expression deconvolution to infer the hierarchy compositions of HCC from bulk transcriptomes. We used multiple scRNAseq-based deconvolution methods and identified ssGSEA as the highest-performing approach among 1,013 hepatocellular carcinoma patients. Patients were clustered into two distinct groups based on the composition of their HCC cell hierarchies: *PGA5*^+^ Hepa group and Proliferate T cell group (**Figure 5a**). In the *PGA5*^+^ Hepa group, both *PGA5*^+^ hepatocyte and *CLDN5*^+^ endothelial exhibited cirrhosis-like features (**Figure 2c**, **Figure S2f**), *RGS5*^+^ endothelial coexisted with *PGA5*^+^ hepatocyte in multiple sets of bulk transcriptomes in cellular interaction (**Figure S5b**). The results revealed that the patients presented a different hierarchical composition at diagnosis and most of the advanced patients were classified into the *PGA5*^+^ Hepa group, such as *PGA5*^+^ hepatocytes were more enriched in advanced patients (**Figure 5b, Figure S6a-S6b**,Wilcoxon test, p value = 1.2e-06). Strikingly, patients with different hierarchy subtypes had different survival outcomes. We found *PGA5*^+^ Hepa hierarchies were associated with worse outcome (**Figure 5c, Figure 5d**). Moreover, in the clinical records of the GSE14520 bulk transcriptome, patients with liver cancer who had a history of cirrhosis had a significantly shorter survival time than other patients (**Figure 5d**). To further validate this result, we performed Cox regression on the 5 cohorts using abundance of HCC cell subpopulations (**see Method**). *PGA5*^+^ Hepa hierarchies (PC1) were predictive of adverse outcomes (**Figure 5e**). Thus, our analysis revealed that hierarchy composition of *PGA5*^+^ hepatocyte was associated with poor prognosis. In addition, no hierarchy was observed in patients with iCCA (**Figure S6c**).

**Figure 5.**
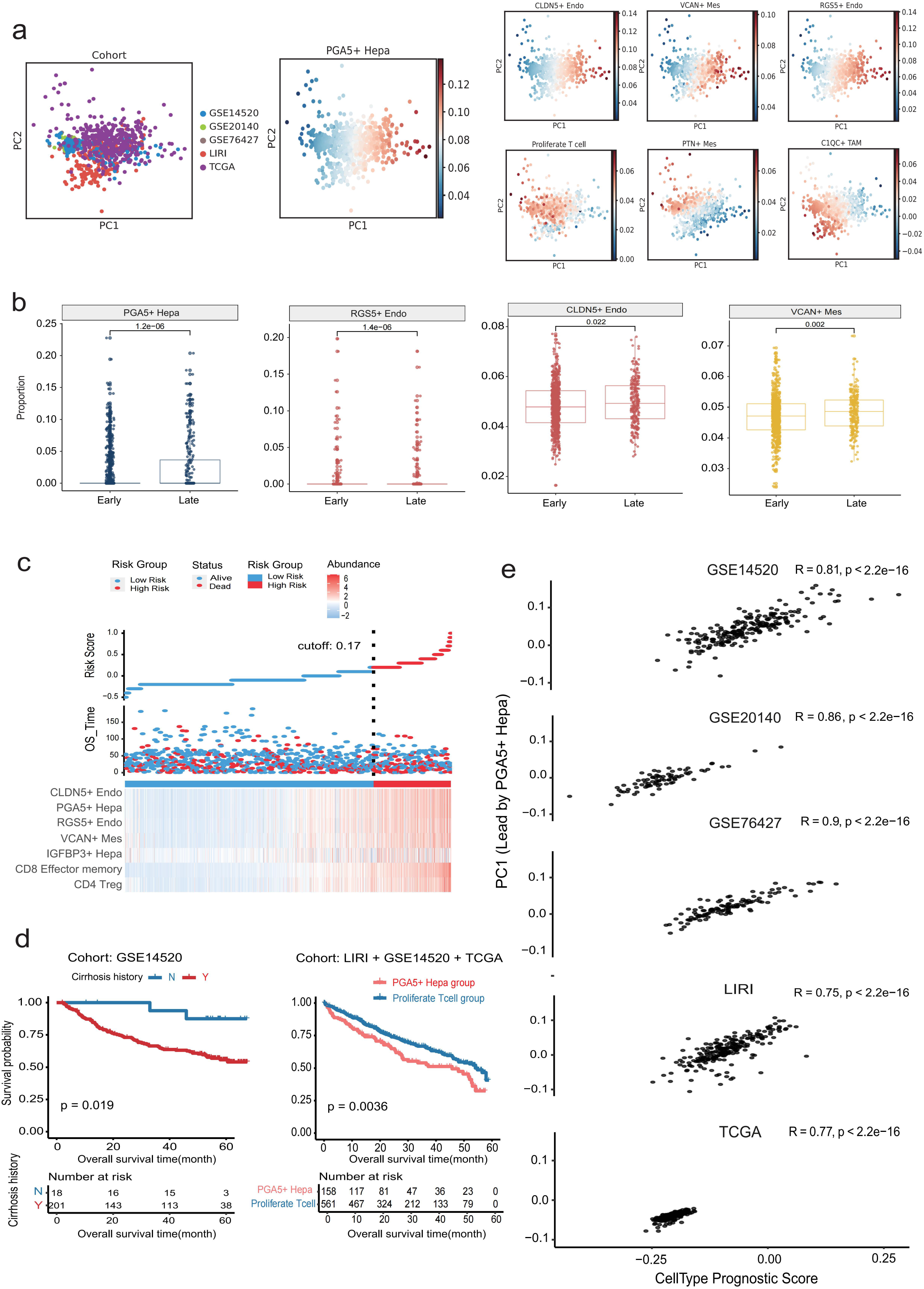
The *PGA5*^+^ Hepa lead axis governs HCC prognosis. **(a)**PCA of 1,013 patients with HCC patients from five cohorts based on the compositions of their cellular hierarchy, which calculated by deconvolution approach using reference signatures from scRNA-seq subpopulations. **(b)** Boxplots show cell proportion of different subtypes in early and late stage patients. **(c)** The risk curve and scatter plot of each sample in the TCGA cohort after realignment via ggrisk algorithm. The heatmap showed distinct expression profiles of seven hierarchies in the high- and low-risk groups. **(d)** Overall survival outcomes of patients which included cirrhosis history within GSE14520 (n=219) (**left**); Overall survival outcomes of patients stratified by PC1 within the LIRI (n = 228), GSE14520 (n=219), and TCGA (n= 371) cohorts (**right**). **(e)** Correlation between a prognostic score trained by regularized cox regression using HCC subtypes abundances with the *PGA5*^+^ Hepa lead proportion axis (PC1) within the five cohorts.

### 3.6. Cirrhosis-like signatures correlate with poor clinical outcome

To further investigate the prognostic value of cirrhosis-like signatures, we screened ten survival associated gene sets (HR>1, risk factor, n=5; HR<1, protective factor, n=5) based on five bulk transcriptomes by using COX regression. We observed that higher enrichment of risk gene sets in cirrhosis and HCC than healthy cell types, while the protective gene sets enriched in healthy samples (**Figure S7a**). Next, we calculated the copy number variation score (**Figure S7b, see Method**) and screened out *KNG1*^+^ hepatocytes that exhibited both prognostic and hypermutation at the cirrhotic stage (**Figure S7c-f**). To identify shared cirrhosis-like signatures associated with survival, we constructed three machine learning models, including XGBoost, random survival forest and LASSO, to seek the most effective prognostic biomarkers (**Figure 6a-6b**). We selected the top 100 shared genes from the cNMF meta-program 2 as inputs to the prediction models (**Figure 1a**). The abundance of these genes were associated with overall survival (**Figure 6c**). After taking variables into the LASSO Cox regression model with minimized lambda (**Figure S8a**), 16 genes (cox.ph>0.05) were selected to build the model(**Figure S8b**). The associations between the expression values of 16 genes and OS were also illustrated in the forestplot, of which only genes such as TTR, TM4SF5 and F2 seemed to be statistically significant (P < 0.05) (**Figure S8b**). We obtained the 10 most significant genes for prognosis using the RSF and XGBoost (**Figure 6a-6b**). Top 5 and top 10 genes were selected to build the model, respectively. We finally obtained better prediction results for the top5 genes (SERPING1, APOC1, HP, CPS1, BLVRB) after XGBoost screening (**Figure 6d)**. Two groups were divided by the abundance of cirrhosis-like signatures after screening. Then Kaplan–Meier survival curve showed that the low expression group had better overall survival, while high expression group had worse prognosis (**Figure 6e**). Furthermore, the cirrhosis-like signature was an independent risk factor in multivariate Cox regression analysis (**Figure 6f**). At the same time, we found that the *PGA5*^+^ subpopulation in HCC could be classified into two subpopulations based on expression of KNG1(**Figure S8c**). Liu et al. found that prognostic models constructed from KNG1 and other genes obtained by differential analysis could devide HCC patients into two different risk groups (2.46 years vs. 6.73 years, p<0.001)(49). We used DEGs including KNG1, TTR, VTN, APOA1, APOC3, and TF**(Figure S8d)**. Survival analysis of four cohorts revealed significant prognostic difference **(Figure S8e)**. Taken together, these findings suggested that the cirrhosis-like signatures were closely associated with clinical outcomes of HCC patients.

**Figure 6.**
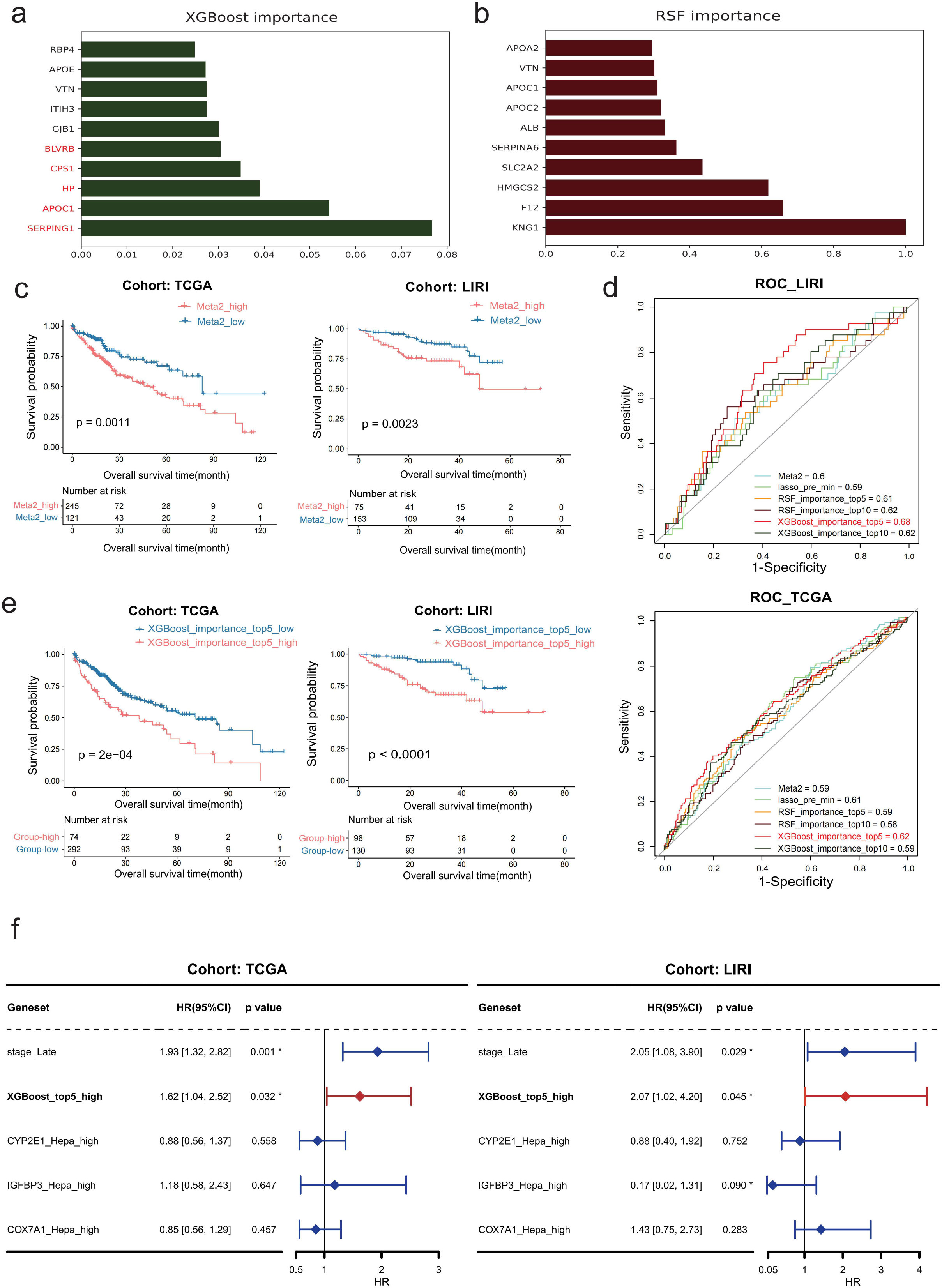
The cirrhosis-like signatures were correlated with poor prognosis. **(a)** Bar plot shows the top 10 important signatures in XGBoost. **(b)** Bar plot shows the top 10 important signatures in RSF. **(c)** Kaplan-Meier curves for overall survival of 366 cases in the TCGA cohort (**left**) and 228 cases in the LIRI cohort (right) stratified by two groups defined by meta-program 2. The p value was calculated by the log-rank test. **(d)** Comparison of multi-class AUCs of three individual machine learning models in the LIRI and TCGA cohorts. **(e)** Kaplan-Meier curves for overall survival of HCC patients in the two cohorts based on two groups defined by “XGBoost_importance_top5” gene set. **(f)** Multivariable analysis of cirrhosis-like signatures in TCGA and LIRI dataset.

## 4. Discussion

Single-cell transcriptome technology could help us understand the heterogeneity of cell types in intertumor, intra-tumor, and precancerous tissue, as well as assist us in exploring the inflammatory and tumor microenvironment. Before that, this technology has been applied to solid tumors, such as the HCC oncofetal characterization(19, 20), and phenotype alignment in colorectal cancer and autologous liver(27). HCC and iCCA are the two most common histologic subtypes. These two molecular subtypes share similar tumor biology and key drivers(50). Cirrhosis, HCC and iCCA are characterized by genomic heterogeneity due to different etiologies(51). Cirrhosis is widely considered as the major risk factor for liver cancer. These studies suggested that similar characteristics of cirrhosis and liver cancer might provide new biomarkers for cancer prognosis. Here, we designed a series of analyses that integrating single-cell transcriptomes and bulk transcriptomes to screen similar subpopulations between cirrhosis and liver cancer, and eventually obtained more accurate predictions of potential markers. In this study, *PGA5*^+^ Hepatocytes in HCC and *KNG1*^+^ Hepatocytes in cirrhosis exhibited similar characteristics at the level of transcriptional programs, GRN, and cellular interactions.

Several limitations should be noted. The integration results in this study could not reveal the molecular mechanism of oncogenesis. The main significance of this work is to establish a data analysis framework to explore molecular similarities between liver cancer and cirrhosis by combining analysis of single cell transcriptome and bulk transcriptomes data. Our study should be further validated in more single cell transcriptomic datasets of liver cirrhosis.

## Contributors

M.T., S.L and L.G.: data interpretation, statistical methods, data analysis and manuscript revision; J.T., Z.Z., C.L, H.H., and Y.L,: manuscript revision; J.H. and M.T.: performed the review and supervised the study.

## Data Sharing Statement

Processed single transcriptome analyzed this study can be downloaded from GEO with the accession number of GSE151530 and GSE136103.

---

**Fig.S1 Cell annotation and correlation analysis.**

**(a)** Uniform manifold approximation and projection (UMAP) embedding of liver cancer and cirrhosis colored by patient ID.

**(b)** UMAP embeddings of single-cell profiles (dots) from individual samples from healthy (**top**), cirrhosis (**middle**), and liver cancer (**bottom**) colored by the expression of signature genes of different cell types.

**(c)** Dot plots of signature genes of different cell types.

**(d)** Heatmap shows the sample preference of each cell type estimated by ratio of observed to expected cell numbers (Ro/e).

**(e)** Boxplots showing that the abundance of each cell type among different samples. (Wilcoxon test, *p < 0.05; **p < 0.01; ***p < 0.001).

(**f, g**) Pearson correlation between average gene expression profiles of subpopulations.

**Fig.S2 Similarity analysis in endothelial cells, MPs and cholangiocytes.**

**(a)** UMAP visualization of cholangiocytes colored by subpopulations among different samples.

**(b)** Sample preference of each subpopulation estimated by the STARTRAC-dist index.

**(c)** Genes screened by the dispersion of genes and the weighting of each cholangiocyte cell subset (**left**). Circle plot showing the phenotypic similarities of various subpopulations of cholangiocytes in three samples (**right**).

**(d)** Umap plot showing the expression level of S100P in malignant cells. Scatterplot of S100P and SPP1 expression.

**(e)** Heatmap shows connectivity (**left**) between cirrhosis-like features enriched cholangiocyte subtypes within iCCA. Jaccard similarity (**right**) of top 100 markers from cirrhosis cholangiocyte subtypes (y axis) with top100 markers of iCCA cholangiocyte subtypes (x axis).

**(f)** Heatmap shows the ratio of observed to expected cell numbers (Ro/e) of each clustering of Endothelial (MPs) cells from different sample origins (**top**). Connectivity (**bottom**) between HCC (x axis) and cirrhosis enriched Endothelial (MPs) clusters (y axis).

**Fig.S3 Stability, error and power in selection of programs in consensus NMF.**

**(a)** Estimated stability (blue, left y axis) and error (red, right y axis) in the cNMF solution learned with different numbers of programs (k, x axis) for hepatocyte (left, range=5~10; right, range=10~50).

**(b)** Heatmap showing correlation of all programs derived from cNMF analysis of HCC and cirrhosis (left, k=9; right, k=49).

**(c)** t-SNE plot of expression levels of selected genes in different clusters indicated by the coloured oval corresponding to **Figure 2f**.

**Fig.S4 TF expression analysis in hepatocyte and cholangiocyte subtypes.**

**(a)** Schematic illustrating the findings of **Figure 3–4**.

**(b)** Endpoint of *PGA5*^+^ (*KNG1*^+^) hepatocyte in the HCC and cirrhosis for Palantir (**top**). Diffusion map of hepatocyte subpopulations based on Palantir (**middle**). The Diffusion map shows TFs expression in different hepatocyte subpopulations (**bottom**).

**(c)** Dotplot shows z-score and RSS of TF regulon (culculated by PySCENIC) among hepatocyte (cholangiocyte) subtypes in different samples (RSS>0.25 were selected for display).

**(d)** Heatmap showing the activity of the top 5 SCENIC-inferred master regulons of nine subpopulations of cholangiocytes.

**(e)** Correlation of binary regulon activity percentage between each cholangiocyte subpopulations and the correlation of p value < 0.001 was shown.

**(f)** Endpoint of *CLDN10*^+^ cholangiocyte in the cirrhosis sample and iCCA for Palantir (**left**). Diffusion map of cholangiocyte subpopulations based on Palantir (**middle**). The Diffusion map shows FOXA3, CEBPD, MEOX1 and NFE2L1 in different cholangiocyte subpopulations (**right**).

**Fig.S5 Cell-cell interaction in hepatocyte and cholangiocyte subtypes.**

**(a)** Number of significant ligand-receptor pairs from *KNG1^+^/PGA5*^+^ hepatocyte (HCC) and *CLDN10*^+^ cholangiocyte (iCCA) to other cells in different sample origin. The edge width is proportional to the indicated number of ligand-receptor pairs.

**(b)** Signatures were extracted from each subtype in scRNA-seq data and were used to calculate subtype proportion in bulk data of HCC (n=5) and iCCA (n=5) by CIBERSORTx. Correlation between subtype proportion in each bulk data is calculated and the abundance is presented in the scatter plots. (R > 0.25: positive correlation, R < −0.25: negative correlation)

**(c)** L-R analysis between hepatocyte (receiver) and T cell (sender) in cirrhosis and HCC samples by NicheNet.

**(d)** The heatmap shows the expression intensity of incoming and outgoing in the intrahepatic cholangiocarcinoma subtypes.

**(e)** The outgoing interaction strength and incoming interaction strength of each cell type in different sample origins. Dot size is proportional to the number of inferred links (both outgoing and incoming) associated with each cell group.

**Fig.S6 Prognosis analysis of HCC data and PCA of iCCA patients.**

**(a)** Boxplots show cell proportion of different subtypes in patients with cirrhosis history.

**(b)** Kaplan-Meier curves for overall survival of HCC patients at early and late stage.

**(c)** PCA of 184 patients with iCCA patients from five cohorts based on the composition of their cellular hierarchy.

**Fig.S7 Prognostic signature enrichment analysis of cirrhosis-like signatures.**

**(a)** Heatmap shows the enrichment score of cell types from different samples among risk (**left**) and protect (**right**) signatures that found in bulk transcriptome by GSVA.

**(b)** UMAP of cirrhosis subtypes by infercnvpy.

(**c, d**) Dotplot shows the extent of copy number variation and prognostic risk/protect factor enrichment in each cell type of the cirrhosis samples, and Dot size represents the significance of the prognostic protect factor enrichment −Log10 (p value).

(**e, f**) Dotplot shows the extent of copy number variation and prognostic risk/protect factor enrichment in each subpopulation of the cirrhosis samples, and Dot size represents the significance of the prognostic protect factor enrichment −Log10 (p value).

**Fig.S8 LASSO regression analysis and survival analysis.**

**(a)** Partial likelihood deviance was revealed by the LASSO regression model. The vertical dotted lines were drawn at the optimal values by using the minimum criteria and 1-SE criteria.

**(b)** The forest plot of the associations between 16 prognostic genes (cox.ph>0.05) and OS in the TCGA cohort. The HR, 95% CI and P value were determined by univariate Cox regression analysis.

**(c)** Violin plot of KNG1 expression in *KNG1*^+^ hepatocyte within cirrhosis and *PGA5*^+^ hepatocyte within HCC.

**(d)** Scatterplot of DEGs between two groups of *PGA5*^+^ hepatocyte.

**(e)** Kaplan-Meier curves for overall survival of HCC patients with high expression of cirrhosis-like and low expression of cirrhosis-like signature in five cohorts.

